# Combined salinity and acidity stressors alter *Daphnia magna* population growth and structure under severe absence of photoperiod

**DOI:** 10.1101/2019.12.11.872630

**Authors:** Mouhammad Shadi Khudr, Samuel Alexander Purkiss, Reinmar Hager

## Abstract

Although natural and anthropogenic influences affect freshwater ecosystems globally at unprecedented levels, the effects of co-occurring physico-chemical stress on zooplankton phenotypic plasticity under extreme conditions remain understudied.

We exposed a laboratory-raised clonal population of *Daphnia magna* to different stress levels of acidity and salinity undergoing complete constant light over 30 days. Overall, population size and age structure at day 10 considerably differed between specific stress contexts. All populations expanded compared to the starting population on day 1. On day 30, overall, population size increased but showed significant differences between treatment groups. Surprisingly, *Daphnia* performed better under combined stress of salinity and acidity than under acidity alone as the extra salinity in the medium may have counterbalanced sodium loss caused by lower pH. Our results reveal a considerable degree of differential reproductive and ontogenetic plasticity in response to combined stressors under disrupted photoperiod. Exposure to constant light led to increased population size, which may be a result of supercharged ion regulation that enables zooplankton to survive better under specific levels of extreme environmental change and adverse chemical stress. Our findings merit further molecular investigation of phenotypic plasticity of the congeners across severe combined stress conditions.

## Introduction

### Co-occurring chemical stressors

Natural and anthropogenic influences on aquatic ecosystems have increased the effects and occurrences of salinisation and acidification of freshwater ecosystems in a number of ways including seawater intrusions, mining, urbanisation and agricultural as well as industrial processes^1–5^. Daphnids are keystone cladoceran zooplankton and important models in the study of environmental stress and biodiversity in aquatic ecosystems^6^. Daphnids produce genetically identical daughters under temperate conditions through parthenogenesis. The genome of *Daphnia* may express different phenotypes in reaction to different environmental conditions; a phenomenon referred to as phenotypic plasticity^7^. Rapid responses to environmental stressors, such as ionic and osmotic regulation^8^, which may be mediated by transgenerational epigenetic effects^9–10^, are associated with high energetic costs^2,11^, and may enable *Daphnia* to occupy a wider niche range compared to other freshwater inhabitants^12^. However, low pH levels have been shown to reduce zooplankton species richness and alter the structure of cladoceran populations^13,14^, leading to diminished survival and growth of *D. magna* below pH 5^15^. For instance, *Daphnia* species are less abundant in acidified lakes compared to other arthropods (*e.g*., calanoid copepods and insects), which may, therefore, dominate the ecosystem instead of *Daphnia^2,16^*.

Like acidification, salinisation of aquatic habitats presents a key stressor of aquatic life. Some species, such as *Daphnia exilis* (Herrick) and *Daphnia pulex* (Leydig), can maintain a positive and constant osmolality difference with the environment at salinities up to 6-8 gL^-1^ via osmoregulation^12^. In particular, *D. magna* (Straus) is a generalist whose niche is defined by a much higher range of salinity tolerance than other aquatic *Daphnia* species. Populations of *D. magna* have been documented to inhabit both fresh and slightly salty waters^17^, with strong evidence of their ability to adjust ion regulation (osmoregulation and osmoconformance) to cope with various levels of salinity^18^. This spans fresh to brackish waters, up to 10 gL^-1^ salinity^19^, with the upper limit of their tolerance depending on population life history and within-species genetic variation^17,20, 21^.

A further ecological challenge arises when changes in salinity may become coupled with water acidification^22^ due to natural and anthropogenic causes leading to increasing levels of salinity and varying levels of acidity in many zooplankton habitats and thus increasingly creating adverse conditions for these organisms^22^. Osmoregulation in *Daphnia* is pH dependent as low pH is thought to inhibit the uptake and increase the loss of sodium much as it does in freshwater fish^21,22^. The negative effects of increased salinisation and acidification on freshwater zooplankton result from impairing the organism’s osmoregulatory system; a compromised ion-regulation can lead to higher sensitivity to low pH, and the latter may alter tolerance to lethal and sub-lethal levels of salinity and *vice versa^22^*. This highlights the need to investigate the effects of co-occurring physical and chemical stressors on keystone species, such as *D. magna,* in freshwater ecosystems^23^.

### Lack of photoperiod as a novel environmental challenge

The duration and intensity of light exposure can have a strong influence on many organisms and research on zooplankton has studied effects on diapause^24,25^; egg hatching^26^, circadian clock, diel vertical migration^27^ and ontogenesis^28^ in response to photoperiod and light quantity^29^. Somewhat surprisingly then, investigations of the effect of exposure to elevated light at night, constant light, or lack of light, on plastic responses of zooplankton in general, and *D. magna* in particular, have been overlooked for decades except for the area of diapause and sexual morph production^25^, and alteration of phototaxis^30^. The negative effects of too little or too much light extend beyond the direct effect on circadian rhythms. Exposure to complete darkness for 7 days has been reported to significantly reduced the survival of *Daphnia parvula* compared to exposure to 14:10 light/dark cycle by a mechanism that is not yet completely understood^31^. On the other end of the spectrum, Meyer & Sullivan (2013) demonstrated that ecological light pollution through increased night lighting modifies community composition, structure, and characteristics and thus ecosystem functioning via alteration of nutrient exchange and aquatic-terrestrial fluxes of invertebrates^32^. These altered fluxes may have cascading effects through riparian and aquatic food webs^33,34^. However, although a number of studies endeavoured to shed light on the effects of light at night on the physiology, behaviour, growth and development, and fitness of a range of aquatic organisms^34–37^, the effect of lack of photoperiod on the phenotype of keystone zooplankton is surprisingly poorly studied. This also includes the effects of existing in waters trapped underground or in caves following storms, tsunamis, and floods^38–41^, or in areas close to the earth’s polar circles^36,42^ that experience both extended periods of Siberian and Nordic nights or midnight sun (*e.g.,*^43^), while being subject to natural and man-made environmental changes. To date, daphnid responses to the implications of ecosystem vulnerability to salinisation and acidification subject to disrupted photoperiod remains largely unexplored.

These patterns and types of complex physico-chemical stress are predicted to lead to a reduction in biodiversity, especially when accompanied by upsetting the thermal regimes in aquatic ecosystems because of climate change and environment modification *(e.g.,* arboreal and riparian vegetation alteration)^34,44,45^. There is a significant gap in current knowledge about how aquatic organisms in general, and zooplankton of economic importance in particular, would respond, in terms of reproductions and development, to the absence of photoperiod, let alone the combination of such physical stress with composite chemical stress (salinisation and acidification of water).

In this work, we aimed to investigate phenotypic plasticity of a clone of *D. magna* under severe conditions of complex abiotic stress as we sought to answer the following main question: How do daphnid reproductive success and age structure differ across varying levels of salinity and acidity, and their combinations, over a 30-day period of exposure to constant light?

## Materials and Methods

### Model organism

The *D. magna* (Straus) clonal population used in this experiment was established from a single female sampled from a population purchased from Sciento©, Manchester UK. The resulting clonal population was reared in a growth chamber, at the Faculty of Biology, Medicine and Health, The University of Manchester, under a 16h:8h light/dark cycle at 23°C. Under these conditions, daphnids reproduce asexually by parthenogenesis resulting in a female-only population. The ability to endure a range of abiotic stressors or sensitivity to contaminants made daphnids a key model to study tolerance^2,46^, phenotypic plasticity^47^, and toxicity in aquatic environments^48^. Feeding occurred every two days by injecting a diet mix into the medium. The diet consisted of 1ml of ‘Allinson dried active yeast’ *(Saccharomyces cerevisiae),* purchased from a local store, and 2ml of alga *Scenedesmus quadricauda* (sourced from Sciento ©, Manchester UK), maintained and quantified in the lab, in Artificial *Daphnia* Medium (ADaM), *sensu* Ebert (2013)^49^, at pH ∼7 and 0.33 gL^-1^ salinity. One litre of ADaM consists of 0.33 g sea salt, 2.3 ml CaCl_2_ x 2H_2_O (117.6 gL^-1^), 2.2 ml NaHCO_3_ (2.52 gL^-1^), and SeO_2_ (0.007 gL^-1^), all dissolved and homogenised in ultra-pure deionised Milli-Q water.

### Experimental design of severe physico-chemical stress

Daphnid population size (total number of daphnids in the beaker at each census, as a proxy of reproductive success) and the proportion of juveniles relative to the total population in the beaker per census (age structure, hereafter) were examined. Adults were identified through observing the brood pouches. The experimental conditions were based on extensive pilot studies during which we established different levels of pH and salinity (chemical stress) that were lethal and sub-lethal to our clone of *D. magna*^47^. The pH treatment levels were 5, 5.5 and 6 and the salinity treatment levels (plus the above-mentioned ADaM salinity) were 1.33, 3.33, and 6.33 gL^-1^. The investigation of salinity and/or acidity stressors, as detailed below, was done in two separate experiments of exposure to physical stress, *i.e.,* lack of photoperiod (constant light).

We used Sigma Aldrich (USA) ‘sea salts’, and distilled acetic acid (Sarson’s ©) procured from a local supplier. To lower the pH in the treatment conditions the acetic acid was added drop-wise to the medium in the beaker and measured continuously using a ‘Mettler Toledo™ FE20 FiveEasy™ benchtop pH meter’ until the desired pH was reached. Here, acetic acid provided a practical way of acidifying the volume of water in question. Acetic acid can have extensive effects on the metabolism of living organisms and it can be found in nature and also close to anthropogenic activity such as bioethanol industry^50^, pond preparation^51^, as a product of anaerobic activity associated with biological decay^52^, waste decomposition and processing^53^, biodegradation of petroleum and hydrocarbon contaminants^54,55^, ocean water, oilfield waters, and both cloud and rain waters^56–60^. Furthermore, adjusting water pH using vinegar is applied in studies on the effects of complex natural organic acids on larval stages of animals *(e.g.,* amphibians^61^). Additionally, the application of vinegar is commonplace in fish farming to neutralise water alkalinity or increase water acidity, owing to proton donation into the aqueous solution 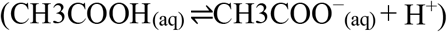. The salinity conditions were achieved by dissolving the correct weight of sea salts in the beaker using a magnetic stirrer and salinity levels were regularly measured using a conductive probe. All experimental beakers were kept in a dedicated growth chamber at 23°C under constant light (24h) regime (lack of photoperiod).

We randomly selected 329 age-synchronised early instars from the stock clone culture reared in the above-mentioned optimal conditions for 20 generations, and allocated from this population 7 instars to 9 different treatment groups (with 5 replicates each) plus a control (the standard ADaM under 24h:0h light/dark cycle, salinity of 0.33 gL^-1^, and pH = ∼7). The treatment groups were as follows: *Salinity1, Salinity2, Salinity3, Acidity1, Acidity2, Acidity3, Salinity1 & Acidity1*, *Salinity2 & Acidity2*, *Salinity3 & Acidity3*). Our pilot study showed that survival was extremely low under such severe experimental conditions as the exposed *Daphnia* clone was not trained/acclimated prior to the exposure. This was done to imitate abrupt as well as extreme environmental change. as provision of food every other day irrespective of animal densities was purposed to take the environmental challenge of the complex physico-chemical stress to the utmost degree beyond the suboptimal, in test of the plasticity of a proportion of the daphnid clone that would manage to wedge, in the Darwinian sense, through adversity into survival and possible subsequent replication.

By Day 10 of the experiment, due to high mortality under severe conditions, only 26 beakers contained live populations from the following treatment groups: *Salinity1* [1.33 gL^-1^, 3 replicates], *Salinity2* [3.33 gL^-1^, 3 replicates], *Salinity3* [6.33 gL^-1^,2 replicates], *Acidity1* [pH 6, 3 replicates], *Acidity2* [pH 5.5, 3 replicates], *Acidity3* [pH 5, 2 replicates], *Salinity1 & Acidity1* [3 replicates], *Salinity2 & Acidity2* [3 replicates], and *Salinity3 & Acidity3* [2 replicates]) plus the control. The beakers (12.5 cm x 10 cm x 10 cm) were cuboid and transparent, each filled with 600 ml of adjusted ADaM. We randomised the positions of the beakers within the growth chamber upon every feeding time. To refresh ADaM, a 75% media change and calibration were conducted weekly because refreshing more often would risk making the severe conditions more hospitable.

Daylight LED bulbs (2800 K Warmwhite 230VAC [NorthLight] purchased from Clas Ohlson ©, Manchester, UK) were used and the light in the growth chamber was surrounding all sides except the base of each beaker, which was placed on a white base. The bulb is a daylight mimic emitting warm soft yellowish light of more than 1100 lumens, which is equivalent to a typical overcast midday in the UK and northern Europe; see also Supplementary Information *Note l*.

### Analysis of *Daphnia* traits

All statistical analyses were conducted in R^62^ via RStudio^63^. The data were collected on Day 10 and Day 30. A generalised mixed effects model (GLMM), family ‘Poisson’, model optimiser ‘bobyqa’^64^, R packages ‘lme4’^65^, was applied to analyse daphnid population size (as defined above) throughout the experiment duration (GLMM 1). There were two censuses. The census (count day) was randomised in the model. This was followed by a posteriori test (Tukey), R package ‘multcomp’^66^, to examine multiple pairwise comparisons of the stressors in question. The explanatory variable (fixed effects) was stressor (salinity, acidity, salinity & acidity) under 24h light comprising the following levels: baseline = the control, *Salinity1* was 1.33 gL^-1^, *Salinity2* was 3.33 gL^-1^, and *Salinity3* was 6.33 gL^-1^; *Acidity1* was pH 6, *Acidity2* was pH 5.5, and *Acidity3* was pH 5; *Salinityl & Acidityl*, *Salinity2 & Acidity2*, and *Salinity3 & Acidity3*. Due to high mortality over time, several treatment combinations and/or replicates were omitted from the analysis. In other words, treatments/replicates without *Daphnia* on Day10 were excluded.

Another model (GLMM 2) was used to analyse age structure per census (as defined above) with the same set up as aforementioned. Besides the model summaries, the main effects of both models are respectively shown via analysis of deviance by using the Anova (Type II) command in R, package ‘car’ ^67^. We note that conducting single analyses of acidity and salinity produced the same pattern of the results outcome retrieved from the approach specified above and thus the single stressor analyses are not presented herein.

## Results and Discussion

### Lack of photoperiod (constant light, 24h), acidity and/or salinity

#### Daphnia population size

Under constant light, 24h, *Daphnia* population size was significantly affected by overall physico-chemical stress treatment (*X*^2^_(9,39)_ = 1117.3, P <0.0001), and specifically by each of the stress levels as follows: *Salinity1* (P < 0.0001), *Salinity2* (P < 0.0001), *Salinity3* (P = 0.03), *Acidity1* (P < 0.0001), *Acidity2* (P < 0.0001), Acidity3 (P < 0.0001), *Salinity1 & Acidity1* (P < 0.0001), *Salinity2 & Acidity2* (P = 0.047), *Salinity3 & Acidity3* (P < 0.0001), (Fig 1); there were 16 daphnids in the control on Day 10 that increased to 328 daphnids on Day 30 (note for reference that under ideal culture conditions of 16:8 light/dark cycle and optimal ADaM the *Daphnia* clone reached from a starting population of 7 instars a population of 35 daphnids on Day 10 increasing to 119 daphnids on Day 30), (Table 1) and Supplementary Information (Tables S1-S3).

**Fig 1.**
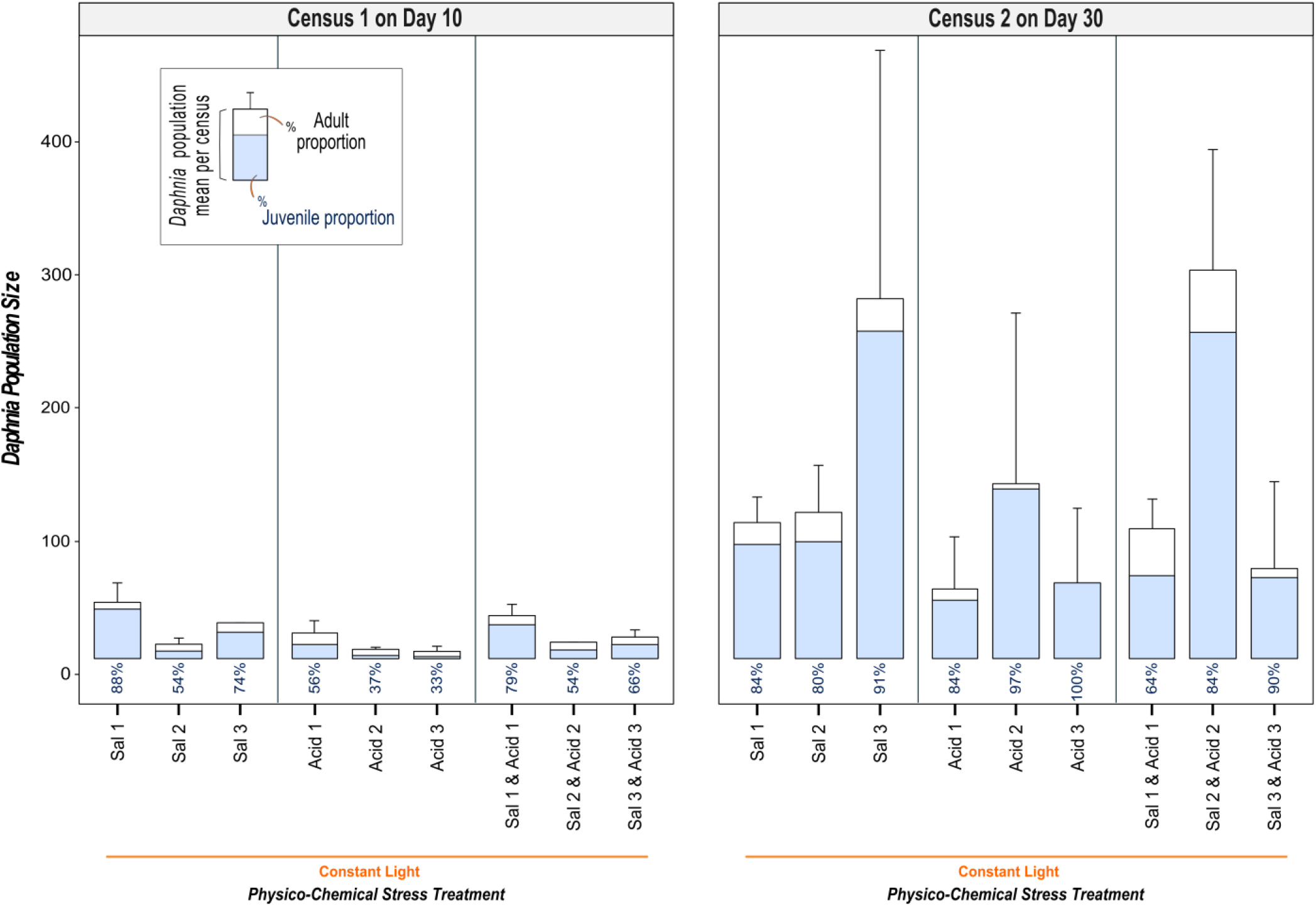
*Daphnia* population sizes and age structure under chemical stress in the absence of photoperiod (constant light). The bars illustrate the changes in *Daphnia* population (mean of total daphnid numbers in the beaker per treatment) ± SEM. The bar parts display the rounded proportions of juveniles against adults relative to the beaker population *(Daphnia* age structure). There were two counts on Day 10 and Day 30. The conditions were as follows: constant light (24h day:0h night), *Salinityl* [1.33 gL^-1^, 3 replicates], *Salinity2* [3.33 gL^-1^, 3 replicates], *Salinity3* [6.33 gL^-1^,2 replicates], *Acidityl* [pH 6, 3 replicates], *Acidity2* [pH 5.5, 3 replicates], *Acidity3* [pH 5, 2 replicates], *Salinityl & Acidityl* [3 replicates], *Salinity2 & Acidity2* [3 replicates], and *Salinity3 & Acidity3* [2 replicates]); Sal = Salinity, Acid = Acidity, Sal & Acid = Salinity & Acidity. The clonal daphnid population in the control (the standard ADaM under 24h:0h light/dark cycle, salinity of 0.33 gL^-1^, and pH = ∼7) was of 16 daphnids on Day 10 and 328 daphnids on Day 30.

**Table 1.** Matrix of multiple pairwise comparisons, *Daphnia* population size under physico-chemical stress. Stress contrasts of the effects of salinity, acidity, and their combination in constant light are presented in a matrix as per the results of the posteriori Tukey test, following GLMM1 (testing *Daphnia* population size [total numbers in the beaker] as specified in the main text Methods). Ctrl (the control, [constant light, 24h day:0h night], salinity of 0.33 gL^-1^, and pH = ∼7), *Salinity1* = 1.33 gL^-1^, *Salinity2* = 3.33 gL^-1^, and *Salinity3* = 6.33 gL^-1^; *Acidity1* (pH = 6), *Acidity2* (pH = 5.5), and *Acidity3* (pH = 5). Only significant results are shown; *ns* refers to non-significance; Sal = Salinity, Acid = Acidity, Sal & Acid = Salinity & Acidity, Light only = constant light without chemical stress, comparable paired labels have matching colour-backgrounds, the orange rectangle label is comparable with all the other labels, the grey rectangles are filling blocks.

|  |  |  |  |  |  |  |  |  |  |
| --- | --- | --- | --- | --- | --- | --- | --- | --- | --- |
|  |  | *Acid3* | P<0.001 | *ns* | P<0.001 |  |  |  |  |
|  |  | *Acid2* | P<0.001 | P<0.001 |  |  |  |  |  |
|  |  | *Acid1* | P<0.001 |  |  |  |  |  |  |
| *Sal2* | *Sal1* |  | Light only<br>(Control) | *Acid1* | *Acid2* | *Acid3* | *Sal1 &*<br>*Acid1* | *Sal2 &*<br>*Acid2* | *Sal3 &*<br>*Acid3* |
| *ns* |  | *Sal1* | P<0.001 | P<0.001 | *ns* | P<0.001 | *ns* | P<0.001 | P<0.001 |
|  | *ns* | *Sal2* | P<0.001 | P<0.001 | *ns* | P<0.001 | *ns* | P<0.001 | P<0.001 |
| P<0.001 | P<0.001 | *Sal3* | *ns* | P<0.001 | P<0.001 | P<0.001 | P<0.001 | *ns* | P<0.001 |
|  |  | *Sal1 &*<br>*Acid1* | P<0.001 | P<0.001 | *ns* | P<0.001 |  |  |  |
|  |  | *Sal2 &*<br>*Acid2* | *ns* | P<0.001 | P<0.001 | P<0.001 | P<0.001 |  |  |
|  |  | *Sal3 &*<br>*Acid3* | P<0.001 | *ns* | P<0.001 | *ns* | P<0.001 | P<0.001 |  |

Compared to Day 1, overall population sizes were considerably larger on Day 10 under stress combinations; notable examples of average percentage of increase/decrease were as follows: *Salinity1* (495% larger population), *Salinity3* (279% larger); *Acidity1* (176% larger), *Acidity2* (5% smaller); *Salinity1 & Acidity1* (362% larger); *Salinity2 & Acidity2* (67% larger), (Fig 1) and Supplementary Information (Table S3). Comparing population sizes on Day 30 to Day 10 reveals that daphnid population size dramatically increased over time with a variable magnitude contingent on the population ability to respond to different stress scenarios. The highest increases displayed under combined stress was *Salinity2 & Acidity2* (2400% larger), followed by *Acidity2* (1865% larger), *Acidity3* (1030% larger) and *Salinity2* (965% larger). By contrast, the smallest increases were seen under *Salinity1* (144% larger) and *Acidity1* (171% larger). The highest level of combined stress, *Salinity3 & Acidity3,* corresponded with a remarkable 352% larger population size. Further, compared to the population at Day 1, the maximal population size, by Day 30, was reached in the combined stress *(Salinity2 & Acidity2’)* (4067% larger), and (3764% larger) in *Salinity3,* while the minimal population increase, by Day 30, was observed under *Acidity1* (648% larger) then *Salinity3 & Acidity3* (857% larger), Supplementary Information (Table S3).

In the short run, acidity stress had a clear negative impact on *Daphnia* population size markedly in *Acidity3* (Fig 1). The poor performance up to Day 10 is attributable to what is known as pH shock^68^ characterised by rapid harmful change in acidity from the stock culture’s semi-neutral pH to the severe experimental acidic conditions. The pH levels of the current study strongly limited *Daphnia* reproductive success and resulted in delayed development to maturity (see age structure below). But, *Daphnia* were able to acclimate to a degree to acidity as the population showed variable resilience over time. By Day 30, a largely juvenile population emphatically recovered in *Acidity3*; *Daphnia* numbers burgeoned in *Acidity2* (mostly juvenile population), whilst the margins of population growth and juvenility were comparatively smaller in *Acidity1* (Fig 1) and Supplementary Information (Table S3). This indicates that the surviving daphnids were more reproductive at pH 5.5 *(Acidity2’)* over time than at pH 6 *(Acidity?)* and pH 5 *(Acidity3’).* Our findings are in line with the observations documented by Weber & Pirow (2009) in *Daphnia pulex*, reporting that tolerance and acclimation, particularly at pH 5.5, can be high and plausibly linked to an induced compensatory active ion transport, owing to *Daphnia’s* ability to lower sodium efflux and increase its influx^2^. It has been shown that in the long run, exposure to pH 5.5 may correspond with better survival and reproduction rates than exposure to pH 5, as values less than 5 can lead to extinction^15^.

Low pH levels are known to reduce survival and growth in *D. magna* due to sodium deficiency^21,22^, a sharp decline in respiratory rates, oxygen uptake depression, and impairment of ion-regulation^15,69^. On the one hand, levels of free-form carbon dioxide (CO_2_) in the surroundings of *Daphnia* concomitantly increase in acidified freshwater, *e.g.,* due to acid spills and acid rain akin to the pH levels tested in this work. At pH 6 the level of free-form CO_2_ is usually more than 60% and at pH 4 this rises to 100%^69^. As such, an external increase in water acidity accompanied by abundant CO_2_ will disturb the internal acid-base balance in *Daphnia;* a degree of haemolymph acidosis that occurs interdependently with decreased CO_2_ diffusion through the gills leading to impeded oxygenation^2^, in accordance with the Bohr effect^69^. In fact, CO_2_-dependent freshwater acidification is a major new challenge for zooplankton; shown by Weiss et al. (2018) to lead to impairment of *Daphnia* detection of predators with consequences for the entire freshwater ecosystem^70^. Those impacts may worsen when ‘acidified’ saltwater intrudes the land by rising sea levels^71,72^.

On the other hand, sodium uptake and osmoregulation in *Daphnia* becomes rapidly destabilised and compromised in acidified water^2,73,74^. The consequence of the above complex interdependence for the functioning of the organism will be energetically costly due to hyper-metabolism caused by increased stress^2^. Daphnids, therefore, may survive the acidity stress dependent on their acidity sensitivity/tolerance^2,74,75^. In sum, daphnid acid-sensitivity and shortage of ambient calcium in acidified water^76^ with subsequent sodium deficit^2^, oxidative stress, increased maintenance under stress^8^, and the resulting costly bioenergetics^2,11^ were the likely reasons behind the poor daphnid performance under the acidity levels of the current work. These findings contribute to our understanding of the damaging effects of low pH levels on aquatic systems that are presently vulnerable to constant change worldwide^5,45^. Like changes in ion concentration in water^5,77^, freshwater acidification is caused by multiple natural and anthropogenic, direct and indirect, short- and long-term causes^78–80^, *e.g.,* acid rain^81^ alongside with organic acids from land catchments and localise acid deposition^78^, pond draining^82^ and drainage waters particularly in areas with soils originated from granite or weathering-resistant mineral aggregates^77,83^, atmospheric and soil deposition of sulphur^78,80,84^, land use and management policies^80,83^, afforestation^80^ and mobilisation of acid anions, via heavy rain, following forest fires and draughts^80,85–87^, use of nitrogen-based fertilisers^88^, melt of snow rich in sulphuric and nitric compounds^81,83,84^, and increased levels of carbon dioxide and pCO_2_ in water^70,89,90^. Indeed, water acidification are projected to lead to alarming ecological consequences with time^70–72,75,89–91^.

For each level of comparison, the differences in population sizes between salinity and acidity stressors were large but then decreased over time; for the contrast *Salinity2 vs Acidity2* the margin of difference between Day 10 (55% larger in salinity) and Day 30 (16% smaller in salinity) is nearly twofold, whilst the difference margin for the contrasts *Salinity1 vs Acidity1* or *Salinity3 vs Acidity3* between Day10 and Day 30 is evidently smaller, Supplementary Information (Table S4). Overall, *Daphnia* population size was notably larger under each level of salinity stress compared to acidity (Day 10: 116%, 55%, and 430%, respectively) and on Day 30 (94%, 16%, and 379%, respectively). Thus, it is clear that salinity was far less damaging to the population than acidity. This relates to halotolerance in *D. magna*^92^ which may have contributed to resilience to the severity of salinity over time in this experiment, but acute and elevated levels of salinity can be detrimental to *Daphnia*^92^ as freshwater salinisation significantly contributes to biodiversity loss and alteration of ecosystem functioning^93,94^.

Both above and below ground freshwater resources are threatened around the globe by elevated levels of salinity within and away from coastal regions due to primary (natural) and secondary (anthropogenic) causes^95^. This includes runoff associated with road-treating salts with consequent groundwater salinisation^93,96^, hydrological alterations of waterways and changes in water quality^4,97,98^, mining and agriculture^4^, and movement of saltwater inland following storm surges, flash floods, rising sea level, and tsunamis^38–41^. For example, the consequences of tsunamis may extend to include the salinisation of coastal/inland fresh surface water, groundwater, and aquifers by means of intrusion, infiltration, and inundation^39,99,100^. It should be noted that our knowledge of both ecosystem and organism responses to alterations of water ionic quality and strength is not comprehensive ^4^’^101^, although increased salt toxicity is predicted to intensify^4,102,103^.

Under normal photoperiods, this can be attributed to osmotic stress and ion cytotoxicity^46^. Osmotic stress occurs when the salt concentration of the environment is higher than the intracellular salt concentration and, as a result, water moves out of the cells. By contrast, cytotoxicity refers to the disruption of the hydrophobic and electrostatic forces that maintain the structure of proteins, inhibiting the normal functioning of enzymes^46^. Our findings show that *Daphnia* was prolific under the highest level of salinity *(Salinity3’)* on Day 30 with a population size ∼169% larger than the observation under *Salinity*1, and ∼161% larger than that under *Salinity2.* This may be due to *Daphnia*’s ability to acclimate to a range of salinity levels by altering their osmotic and ionic regulation when water acidity and salinity levels are sub-lethal^8^, (Fig 1) and Supplementary Information (Table S3)

To our surprise, the effect of the chemical stressors under constant-light stress led to unexpected results. *Daphnia magna* showed the same or greater population growth under combined salinity and acidity. On Day 10, there was greater growth in 1 (*vs Salinity2*) out of 3 scenarios compared to salinity alone; and in 3 out of 3 compared to acidity alone. On Day 30, the population size under combined salinity and acidity was larger in 1 (*vs Salinity2*) out of 3 scenarios compared to salinity, and in 2 out of 3 compared to acidity, (Fig 1) and Supplementary Information (Table S5). In contrast to our prediction, across all treatment groups, population size was the largest in the combined stress *Salinity2 & Acidity2.* In this group, there were on average 291.66 individuals, generated from as little as 12 individuals on Day 10. The value of ∼292 individuals on Day 30 was 165% and 123%, respectively, larger than what was achieved by *D. magna* in *Salinity2* and *Acidity2*. Further, the population size under combined stress, *Salinity2 & Acidity2*, was respectively ∼8%, ∼187%, ∼416%, ∼457%, ∼201%, and ∼335% larger than what were achieved in *Salinity3, Salinity1*, *Acidity3*, *Acidity1*, *Salinity1 & Acidity1*, and *Salinity3 & Acidity3,* respectively; see (Fig 1) and Supplementary Information (Tables S4 and S5). As highlighted above, the uptake of sodium is reduced in acidic conditions as sodium uptake takes place in exchange for hydrogen ions^73^, where the rate of sodium loss in *D. magna* is documented to increase fourfold between pH 7 and pH 3^73^, leading to death by reduced uptake of environmental sodium, especially under low concentrations^73^. This might explain why *D. magna* in *Salinity2 & Acidity2* obviously outperformed the reproductive success in *Salinity1 & Acidity1* and *Salinity3 & Acidity3*. This result could be linked to the capacity of the daphnid genotype to rapidly respond to ecological challenge and preconditioning offspring through adopting different strategies^9,10,104^, ensuring prioritising energy allocation towards reproduction as opposed to growth under stress^11,94^. The combined effect of salinity and acidity under constant light on population sizes in our work was more severe than each of the individual stressors on Day 10. However, by Day 30, the combined stress effect became more than additive for level 3 of salinity or acidity, considerably less than additive for level 2, and almost equal to the total effect of salinity for level 1, (Fig 1) and Supplementary Information (Table S5). This is in line with the findings documented by Zalizniak et al. (2009) who experimented on low pH effect on lethal and sub-lethal salinity tolerance in macro-invertebrates^22^; they found that abundant ambient salinity may offset the negative effect of acidity perhaps through exchanging the excess protons in the haemolymph with sodium. In our study, however, the fact that population size under *Salinity3 & Acidity3* was larger than the population size under *Acidity3*, at the first census by 11% then smaller by the same percentage at the second census, but in both cases smaller than under salinity alone, is partly in accord with the common view that increased acidity should aggravate salinity effects^22^., and becomes more intriguing given the fact that higher sodium concentration in the ambient medium can aid sodium turnover^22^. Overall, our results corroborate the view advocated by Altshuler et al. (2011) that a comprehensive understanding of the combined effects of multiple stressors will require the specific assessment of the interactive effects of these stressors rather than individually examining the effects of each stressor^23^. Furthermore, a causative relationship between physico-chemical stressors may exist as changes in one stressor can lead to changes in other stressors, *e.g.*, freshwater salinisation syndrome^4,77,105^. Hence, one could argue that complex toxic effects of stressor interactions may underlie our results. To date, a comprehensive understanding of the physiological responses to ionic changes (alone and accompanied with pH changes) is still developing^4,103^. Coping with the combined chemical stress that is naturally unfavourable for exponential reproduction can be plausibly associable with a compensatory metabolic response under constant exposure to light through an as yet unknown mechanism. Light pollution effects, in terms of artificial light, sky glow and interference with daylight-night-time patterns, which increasingly affect aquatic ecosystems and ecological interactions^34,106^, are becoming a serious concern associated with urbanisation and industrialised zones with increasing human population around the globe^30,34,107–110^. The far-reaching effects of extended light exposure^34^ can not only lead to changes in vital phototactic behaviours with disruptive impacts on the aquatic system^30,34^, but may also extend to how zooplankton respond to a cocktail of stressors^77,111^; see Supplementary Information *Note 2* for further discussion of *Daphnia* coping mechanisms against complex ecological challenge.

#### Daphnia population age structure

Under lack of photoperiod (constant light, 24h), *Daphnia* juvenile proportions were significantly affected by overall physico-chemical stress treatment (X^2^_(9,36)_ = 119.89, P <0.0001), and specifically *only* by the following stress levels: *Acidity2* (P = 0.035) and *Acidity3* (P < 0.0001), (Fig 1) and Supplementary Information (Tables S6 and S7). There was an abrupt decrease in adult proportion from 67% adults on Day 10 to 0% on Day 30 in *Acidity3,* and a sharp drop from 46% to 20% in *Salinity2*. Interestingly, only in 2 out of all the 10 stress treatments were there more adults than juveniles on Day 10 with respect to *Acidity2* (63% adults) and *Acidity3* (67% adults); this was markedly reversed on Day 30 as significantly more juveniles were recorded then under each of these stress scenarios, (Fig 1) and Supplementary Information (Table S8). Conversely, only two cases of the remainder (7 scenarios out of 10), *Salinity1* and combined stress *Salinity1 & Acidity1*, led to noticeable increases in adults on Day 30 compared to Day 10, (Fig 1) and Supplementary Information (Table S8). Moreover, it is worth pointing out that the proportions of juveniles (84%) and adults (16%), on Day 30 under constant light for *Salinity1*, *Acidity1*, and combined stress *Salinity2 & Acidity2*, were identical despite the very different stress conditions in these treatments, and the respective differences in average population size at Day 30 (∼102 individuals, ∼52 individuals, and ∼292 individuals, respectively). However, overall, there were more adults on Day 10 in the acidity than in the salinity levels. In contrast, the opposite was observed on Day 30 except for *Salinity1 vs Acidity1* where the adult proportions were identical irrespective of the larger population size achieved under *Salinity1,* (Fig 1) and Supplementary Information (Tables S4 and S8).

*Daphnia* showed the greatest resilience under *Salinity2, Salinity3, Acidity2* (pH 5.5), and combined stress *Salinity2 & Acidity2* as the population increased from means of (∼10, ∼27, ∼7, and ∼12 individuals, respectively) on Day 10 to means of (110, 271, 131, and 292, individuals, respectively) on Day 30, with large proportions of juveniles. Having a prolonged juvenile period seemed to help *D. magna* in coping, as a form of resilience. Note that by Day 30 under the highest level of water acidification in *Acidity3* the entire population consisted of juveniles. However, such resilience was less apparent at Day 30 in *Acidity1* and combined stress, *Salinity1 & Acidity1*, as more adults were found there in the population. At any rate, there were more juveniles with delayed maturation when the photoperiod was lacking (constant light), but generally the population on Day 30 was more juvenile under acidity stress than salinity or the combination of both stressors. This corresponds with the findings reported by De Coen & Janssen (2003)^112^ that exposure to sub-lethal chemical stressors might cause a decreased ‘net energy budget’ in *D. magna,* which means that the physiological energetics of the stressed daphnids will be shifted more towards costly compensatory metabolic mechanisms^11^, such as respiration and exoskeleton formation^76^ rather than growth and reproduction. This view is supported by Glover & Wood (2005) who suggested that *D. magna* sensitivity to acidic environments can be explained by a dependency of sodium influx on sodium-proton exchange system and co-occurrent remineralisation of the exoskeleton^74^. Acidic media (with low levels of sodium and calcium), accompanied by exhausted respiration and ion-regulation, may slow metamorphosis, which occurs approximately twice per week in optimal conditions. In turn, this may benefit *D. magna* in mitigating the flux deficit of essential minerals with less allocation of minerals to re-mineralise the exoskeleton. Such a survival strategy of more investment in longevity may be advantageous until conditions become more hospitable^11^. Furthermore, the rates of sodium turnover in *Daphnia* are high when compared to those of other fresh water animals. However, the rate of sodium turnover is related to their small size and greater surface area to volume ratio^73^, which is even larger in juveniles than adults^73^. Moreover, Na^+^ uptake in *D. magna* is ATP-dependent (Na^+^/K^+^ pump), which is different in juveniles compared to adults due to developmentally dependent biochemical differences^113^. The affinity constant for sodium transport is much lower in adults, and juveniles have a lower affinity for sodium, which is counterbalanced by a higher maximum capacity of sodium uptake^113^. It can be argued that delayed maturation is a plastic trait governed by maternal preconditioning of offspring phenotype^9,10,104^, akin to diapause, *i.e.,* resting^28^, which will favour survival rather than reproduction in adverse environmental conditions. This may be beneficial when accompanied by the ability of the preconditioned neonates to alter their ionic and osmotic regulation^8^. In this respect, a delay of reproduction for the advantage of investment in energy accumulation, to be made available for sequent reproduction when the severity of conditions lessens, has been shown as a survival tactic by *Daphnia* on deficient/limited resources^114,115^. However, these survival strategies may be context-dependent, clone-specific^17^ and again affected by transgenerational effects^9,10^. Overall, the excess ambient amount of Na^+^ in combined stress (*Salinity2 & Acidity2* and *Salinity3 & Acidity3*) could have played a vital role in mitigating the negative impact of acidification on exhausting the haemolymph content of sodium^74^. This could also explain the high mortality in acidity alone treatments in our study. Both weak and strong acidification are known to directly and indirectly negatively affect the physiology, fitness, and survival of freshwater biota^70,116^ and thus leading to mortality and biodiversity loss^75,117^, especially for taxa such as *Daphnia* that are sensitive to pH changes^74,75^. Finally, we observed no egg production under any conditions although daphnids are known to produce diapausing eggs during periods of environmental stress induced by natural and anthropogenic factors (e.g.,^118,119^); this further suggests that our results may be reflective of plasticity found in *D. magna*^47,120^.

## Conclusion

This work is the first to demonstrate that exposure to physical stress (prolonged disrupted photoperiod, constant light) combined with chemical stresses (salinity and/or acidity) can differentially alter *Daphnia* population growth and age structure over time, and may underlie a degree of acclimation to severe chemical stress. Our findings are a first step to a more comprehensive understanding of how aquatic organisms adapt to extreme complex conditions under prolonged disruption of photoperiod. Exploration of the metabolomic regulation of uptake and efflux of haemolymph ions needs to be the next step to test the organism’s response and adaptability under combinations of multiple physical and chemical stressors.

## Acknowledgements

We thank Alice de Sampaio Kalkuhl for her help in the maintenance of the system and in data collection.

## Competing interests

The authors have declared that no competing interests exist.

## Data Availability

The data analysed in this work are available at the figshare data repository via the URL: https://figshare.com/s/34d8a98c279b2dc2cb81

## Notes

https://figshare.com/s/34d8a98c279b2dc2cb81

### Competing Interest Statement

The authors have declared no competing interest.

### Summary of Updates

Shortened and updated Discussion

