## Supplementary Information for "Combined salinity and acidity stressors alter *Daphnia magna* population growth and structure under severe absence of photoperiod"

**Running title:** Combined physico-chemical stress affects *Daphnia* fitness

**Table S1. Effects of physico-chemical stress treatment levels on population size under constant light (24h).** Given are the model summary and estimates from the GLMM 1 for each level of stress treatment together with the standard error, z-value and the associated p-value. The model tests, across treatment combinations, *Daphnia* population size (total numbers in the beaker as specified in the main text Methods), Ctrl (the control, absence of photoperiod [constant light, 24h day:0h night], salinity of 0.33 $gL^{-1}$, and pH = ~7), *Salinity1* = 1.33 $gL^{-1}$, *Salinity2* = 3.33 $gL^{-1}$, and *Salinity3* = 6.33 $gL^{-1}$; *Acidity1* (pH = 6), *Acidity2* (pH = 5.5), and *Acidity3* (pH = 5); Sal = Salinity, Acid = Acidity, Sal & Acid = Salinity & Acidity.

| **Fixed effects** | Estimate | Std. Error | z value | Pr(>\|z\|) |
|---|---|---|---|---|
| (Intercept) | 4.72 | 0.7 | 6.7 | **<0.0001** |
| **Physico-chemical stress under 24h light** | | | | |
| *Salinity1* | -0.88 | 0.07 | -12.1 | **<0.0001** |
| *Salinity2* | -1.05 | 0.08 | -13.94 | **<0.0001** |
| *Salinity3* | -0.15 | 0.07 | -2.17 | **0.03** |
| *Acidity1* | -1.57 | 0.09 | -18. | **<0.0001** |
| *Acidity2* | -0.92 | 0.07 | -12.55 | **<0.0001** |
| *Acidity3* | -1.72 | 0.11 | -16.39 | **<0.0001** |
| *Salinity1 & Acidity1* | -0.98 | 0.07 | -13.21 | **<0.0001** |
| *Salinity2 & Acidity2* | -0.13 | 0.06 | -1.99 | **0.047** |
| *Salinity3 & Acidity3* | -1.42 | 0.09 | -15.05 | **<0.0001** |

**Table S2. Stress posteriori contrasts, *Daphnia* population size under salinity, acidity, and their combination in constant light.** Multiple pairwise comparisons (Tukey test), following GLMM 1 (testing *Daphnia* population size [total numbers in the beaker] under combined stress as specified in the main text Methods). Ctrl (the control, absence of photoperiod [constant light, 24h day:0h night], salinity of 0.33 gL$^{-1}$, and pH = ~7), *Salinity1* = 1.33 gL$^{-1}$, *Salinity2* = 3.33 gL$^{-1}$, and *Salinity3* = 6.33 gL$^{-1}$; *Acidity1* (pH = 6), *Acidity2* (pH = 5.5), and *Acidity3* (pH = 5). Only significant results are shown; Sal = Salinity, Acid = Acidity, Sal & Acid = Salinity & Acidity.

| Linear Hypotheses | Estimate | Std. Error | z value | Pr(>\|z\|) |
|---|---|---|---|---|
| *Sal1* ***vs*** Ctrl == 0 | -0.88 | 0.07 | -12.1 | **<0.001** |
| *Sal2* ***vs*** Ctrl == 0 | -1.05 | 0.08 | -13.94 | **<0.001** |
| *Sal3* ***vs*** *Sal1* == 0 | 0.73 | 0.06 | 11.51 | **<0.001** |
| *Sal3* ***vs*** *Sal2* == 0 | 0.9 | 0.067 | 13.54 | **<0.001** |
| *Acid1* ***vs*** Ctrl == 0 | -1.57 | 0.09 | -18.05 | **<0.001** |
| *Acid2* ***vs*** Ctrl == 0 | -0.92 | 0.07 | -12.55 | **<0.001** |
| *Acid3* ***vs*** Ctrl == 0 | -1.72 | 0.11 | -16.34 | **<0.001** |
| *Acid1* ***vs*** *Acid2* == 0 | 0.65 | 0.08 | 7.76 | **<0.001** |
| *Acid2* ***vs*** *Acid3* == 0 | -0.81 | 0.1 | -7.85 | **<0.001** |
| *Acid1* ***vs*** *Sal1* == 0 | -0.69 | 0.08 | -8.3 | **<0.001** |
| *Acid1* ***vs*** *Sal2* == 0 | -0.52 | 0.09 | -6.02 | **<0.001** |
| *Acid1* ***vs*** *Sal3* == 0 | -1.42 | 0.08 | -17.87 | **<0.001** |
| *Acid2* ***vs*** *Sal3* == 0 | -0.77 | 0.06 | -12 | **<0.001** |
| *Acid3* ***vs*** *Sal1* == 0 | -0.85 | 0.1 | -8.28 | **<0.001** |
| *Acid3* ***vs*** *Sal2* == 0 | -0.67 | 0.1 | -6.43 | **<0.001** |
| *Acid3* ***vs*** *Sal3* == 0 | -1.57 | 0.1 | -15.9 | **<0.001** |
| (*Sal1 & Acid1*) ***vs*** Ctrl == 0 | -0.98 | 0.07 | -13.21 | **<0.001** |
| (*Sal3 & Acid3*) ***vs*** Ctrl == 0 | -1.42 | 0.09 | -15.05 | **<0.001** |
| (*Sal2 & Acid2*) ***vs*** *Sal1* == 0 | 0.75 | 0.059 | 12.8 | **<0.001** |
| (*Sal3 & Acid3*) ***vs*** *Sal1* == 0 | -0.55 | 0.09 | -5.98 | **<0.001** |
| (*Sal2 & Acid2*) ***vs*** *Sal2* == 0 | 0.92 | 0.06 | 14.87 | **<0.001** |
| (*Sal3 & Acid3*) ***vs*** *Sal2* == 0 | -0.37 | 0.09 | -3.96 | **<0.001** |
| (*Sal1 & Acid1*) ***vs*** *Sal3* == 0 | -0.83 | 0.07 | -12.74 | **<0.001** |
| (Sal3 & Acid3) ***vs*** Sal3 == 0 | -1.27 | 0.09 | -14.52 | **<0.001** |
| (*Sal1 & Acid1*) ***vs*** *Acid1* == 0 | 0.59 | 0.09 | 6.95 | **<0.001** |
| (*Sal2 & Acid2*) ***vs*** *Acid1* == 0 | 1.44 | 0.08 | 19.03 | **<0.001** |
| (*Sal2 & Acid2*) ***vs*** *Acid2* == 0 | 0.79 | 0.06 | 13.32 | **<0.001** |
| (*Sal3 & Acid3*) ***vs*** *Acid2* == 0 | -0.51 | 0.09 | -5.51 | **<0.001** |
| (*Sal1 & Acid1*) ***vs*** *Acid3* == 0 | 0.74 | 0.1 | 7.19 | **<0.001** |
| (*Sal2 & Acid2*) ***vs*** *Acid3* == 0 | 1.6 | 0.1 | 16.61 | **<0.001** |
| (*Sal2 & Acid2*) ***vs*** (*Sal1 & Acid1*) == 0 | 0.85 | 0.06 | 14.06 | **<0.001** |
| (*Sal3 & Acid3*) ***vs*** (*Sal1 & Acid1*) == 0 | -0.44 | 0.09 | -4.78 | **<0.001** |
| (*Sal3 & Acid3*) ***vs*** (*Sal2 & Acid2*) == 0 | -1.3 | 0.08 | -15.36 | **<0.001** |

**Table S3. *Daphnia* population size differences under salinity, acidity, and their combination in constant light.** The changes (positive or negative) in average population size (‘Pop. size’), mean of total numbers in the beaker as specified in the main text Methods, across all treatment groups are shown in rounded percentage. There were two censuses (Day 10 and Day 30). Ctrl (the control, absence of photoperiod [constant light, 24h day:0h night]), salinity of 0.33 $gL^{-1}$, and pH = ~7), *Salinity1* = 1.33 $gL^{-1}$, *Salinity2* = 3.33 $gL^{-1}$, and *Salinity3* = 6.33 $gL^{-1}$; *Acidity1* (pH = 6), *Acidity2* (pH = 5.5), and *Acidity3* (pH = 5). Sal = Salinity, Acid = Acidity, Sal & Acid = Salinity & Acidity, Photo.p. = Photoperiod, absence of photoperiod (constant light, 24h day:0h night), Pop. = Population, +% = proportionally larger.

| | | Initial Pop. | Pop. size (mean) | % Pop. size expansion or shrinkage | Pop. size (mean) | % Pop. size expansion or shrinkage | Difference in pop. size | % Pop. size expansion or shrinkage | % Pop. size expansion or shrinkage | |
|---|---|---|---|---|---|---|---|---|---|---|
| **Photo.p.** | **Stress Scenario** | **Day 1** | **Day 10** | **Day 10 Compared to Day 1** | **Day 30** | **Day 30 Compared to Day 1** | **between Day 10 and Day 30** | **Day 30 Compared to Day 10** | **On Day 10 Compared to Ctrl** | **On Day 30 Compared to Ctrl** |
| 24D:0N | Ctrl | 7 | 16 | 129% larger | 328 | 4586% larger | 312 | 1950% larger | NA | NA |
| 24D:0N | *Sal1* | 7 | 42 | 495% larger | 102 | 1352% larger | 60 | 144% larger | 160% larger | 69% **smaller** |
| 24D:0N | *Sal2* | 7 | 10 | 48% larger | 110 | 1471% larger | ~100 | 965% larger | 35% **smaller** | 66% **smaller** |
| 24D:0N | *Sal3* | 7 | 27 | 279% larger | 271 | 3764% larger | 244 | 921% larger | 66% larger | 18% **smaller** |
| 24D:0N | *Acid1* | 7 | 19 | 176% larger | 52 | 648% larger | 33 | 171% larger | 21% larger | 84% **smaller** |
| 24D:0N | *Acid2* | 7 | **~7** | 5% **smaller** | 131 | 1771% larger | 124 | 1865% larger | 58% **smaller** | 60% **smaller** |
| 24D:0N | *Acid3* | 7 | 5 | 29% **smaller** | 57 | 707% larger | 52 | 1030% larger | 69% **smaller** | 83% **smaller** |
| 24D:0N | *Sal1 & Acid1* | 7 | 32 | 362% larger | 97 | 1286% larger | 65 | 200% larger | 102% larger | 70% **smaller** |
| 24D:0N | *Sal2 & Acid2* | 7 | 12 | 67% larger | 292 | 4067% larger | 280 | 2400% larger | 27% **smaller** | 11% **smaller** |
| 24D:0N | *Sal3 & Acid3* | 7 | 16 | 129% larger | 67 | 857% larger | 51 | 352% larger | 0% | 261% **smaller** |

**Table S4. Meta-comparisons of *Daphnia* reproductive success under salinity and acidity in constant light.** Mean population size (a proxy for reproductive success) and changes (in rounded percent) in population size in salinity compared to acidity are shown for two censuses (Day 10 and Day 30) across all treatment levels. Photoperiod was absent (constant light, 24h day:0h night). *Salinity1* = 1.33 $gL^{-1}$, *Salinity2* = 3.33 $gL^{-1}$, and *Salinity3* = 6.33 $gL^{-1}$; *Acidity1* (pH = 6), *Acidity2* (pH = 5.5), and *Acidity3* (pH = 5).

| | **Population size (mean) on Day 10** | | | Comparative% increase/decrease |
|---|---|---|---|---|
| Stressor Level | Salinity | Acidity | | *Salinity compared to Acidity* |
| 1 | 41.67 | 19.33 | | 116% **larger** |
| 2 | 10.33 | 6.67 | | 55% **larger** |
| 3 | 26.5 | 5 | | 430% **larger** |
| | **Population size (mean) on Day 30** | | | Comparative% increase/decrease |
| Stressor Level | Salinity | Acidity | | *Salinity compared to Acidity* |
| 1 | 101.66 | 52.33 | | 94% **larger** |
| 2 | 110 | 131 | | 16% smaller |
| 3 | 270.5 | 56.5 | | 379% **larger** |

**Table S5. Meta-comparisons of *Daphnia* reproductive success under salinity, acidity, and their combination in constant light.** Mean population size (a proxy for reproductive success) and changes (in rounded percent) in population size in combined chemical stress (salinity & acidity) compared to salinity or acidity are shown for two censuses (Day 10 and Day 30) across all treatment levels. Photoperiod was absent (constant light, 24h day:0h night). *Salinity1* = 1.33 $gL^{-1}$, *Salinity2* = 3.33 $gL^{-1}$, and *Salinity3* = 6.33 $gL^{-1}$; *Acidity1* (pH = 6), *Acidity2* (pH = 5.5), and *Acidity3* (pH = 5).

| | **Population size (mean) on Day 10** | | | | Comparative % increase/decrease under Salinity & Acidity | |
|---|---|---|---|---|---|---|
| Stressor Level | Salinity | Acidity | Salinity & Acidity | | *compared to* Salinity | *compared to* Acidity |
| 1 | 41.67 | 19.33 | 32.33 | | 22% smaller | 67% **larger** |
| 2 | 10.33 | 6.67 | 11.66 | | 13% **larger** | 75% **larger** |
| 3 | 26.5 | 5 | 10.66 | | 60% smaller | 11% **larger** |
| | **Population size (mean) on Day 30** | | | | Comparative % increase/decrease under Salinity & Acidity | |
| Stressor Level | Salinity | Acidity | Salinity & Acidity | | *compared to* Salinity | *compared to* Acidity |
| 1 | 101.66 | 52.33 | 97 | | 5% smaller | 85% **larger** |
| 2 | 110 | 131 | 291.67 | | 165% **larger** | 123% **larger** |
| 3 | 270.5 | 56.5 | 44.67 | | 83% smaller | 11% smaller |

**Table S6. Effects of physico-chemical stress treatment on age structure in constant light (24h).** Given are the model summery and estimates from the GLMM 2 for each level of stress treatment together with the standard error, z-value and the associated p-value. The model tests the proportions of juveniles relative to the total beaker population as an indicator of *Daphnia* age structure (specified in the main text Methods) across treatment combinations, Ctrl (the control, absence of photoperiod [constant light, 24h day:0h night], salinity of 0.33 $gL^{-1}$, and pH = ~7), *Salinity1* = 1.33 $gL^{-1}$, *Salinity2* = 3.33 $gL^{-1}$, and *Salinity3* = 6.33 $gL^{-1}$; *Acidity1* (pH = 6), *Acidity2* (pH = 5.5), and *Acidity3* (pH = 5); Sal = Salinity, Acid = Acidity, Sal & Acid = Salinity & Acidity.

| **Fixed effects** | Estimate | Std. Error | z value | Pr(>\|z\|) |
|---|---|---|---|---|
| (Intercept) | 4.34 | 0.12 | 36.64 | **<0.0001** |
| **Physico-chemical Stress under 24h light** | | | | |
| *Salinity1* | 0.11 | 0.09 | 1.23 | 0.22 |
| *Salinity2* | -0.14 | 0.09 | -1.47 | 0.14 |
| *Salinity3* | 0.07 | 0.1 | 0.68 | 0.5 |
| *Acidity1* | -0.12 | 0.1 | -1.24 | 0.22 |
| *Acidity2* | -0.21 | 0.1 | -2.11 | **0.035** |
| *Acidity3* | -1.2 | 0.15 | -8.17 | **<0.0001** |
| *Salinity1 & Acidity1* | -0.08 | 0.09 | -0.81 | 0.42 |
| *Salinity2 & Acidity2* | -0.1 | 0.09 | -1.11 | 0.27 |
| *Salinity3 & Acidity3* | 0.01 | 0.1 | 0.1 | 0.92 |

**Table S7. Posteriori contrasts, *Daphnia* age structure under salinity, acidity, and their combination in constant light (24h).** Multiple pairwise posthoc comparisons (Tukey), following GLMM 2 (testing *Daphnia* age structure [proportions of juveniles relative to the total population in the beaker] as described in the main text Methods). Ctrl (the control, photoperiod of 24h light:0h darkness, salinity of 0.33 $gL^{-1}$, and pH = ~7), *Salinity1* = 1.33 $gL^{-1}$, *Salinity2* = 3.33 $gL^{-1}$, and *Salinity3* = 6.33 $gL^{-1}$; *Acidity1* (pH = 6), *Acidity2* (pH = 5.5), and *Acidity3* (pH = 5). Only significant results are shown; Sal = Salinity, Acid = Acidity, Sal & Acid = Salinity & Acidity.

| Linear Hypotheses | Estimate | Std. Error | z value | Pr(>\|z\|) |
|---|---|---|---|---|
| *Acid3* ***vs*** Ctrl == 0 | -1.2 | 0.15 | -8.17 | **<0.01** |
| *Acid3* ***vs*** *Acid1* == 0 | -1.08 | 0.13 | -8.04 | **<0.01** |
| *Acid3* ***vs*** *Acid2* == 0 | -0.99 | 0.13 | -7.33 | **<0.01** |
| *Sal2* ***vs*** *Sal1* == 0 | -0.25 | 0.07 | -3.78 | **<0.01** |
| *Acid1* ***vs*** *Sal1* == 0 | -0.23 | 0.07 | -3.32 | **0.028** |
| *Acid2* ***vs*** *Sal1* == 0 | -0.32 | 0.07 | -4.44 | **<0.01** |
| *Acid3* ***vs*** *Sal1* == 0 | -1.31 | 0.13 | -10.08 | **<0.01** |
| *Acid3* ***vs*** *Sal2* == 0 | -1.06 | 0.13 | -8.01 | **<0.01** |
| *Acid2* ***vs*** *Sal3* == 0 | -0.27 | 0.08 | -3.45 | **0.018** |
| *Acid3* ***vs*** *Sal3* == 0 | -1.26 | 0.134 | -9.42 | **<0.01** |
| (*Sal1 & Acid1*) ***vs*** *Acid3* == 0 | 1.12 | 0.13 | 8.52 | **<0.01** |
| (*Sal2 & Acid2*) ***vs*** *Acid3* == 0 | 1.09 | 0.13 | 8.29 | **<0.01** |
| (*Sal3 & Acid3*) ***vs*** *Acid3* == 0 | 1.21 | 0.13 | 8.95 | **<0.01** |
| (*Sal2 & Acid2*) ***vs*** *Sal1* == 0 | -0.22 | 0.07 | -3.3 | **0.03** |

**Table S8. *Daphnia* age structure under salinity, acidity, and their combination in constant light (24h).** Proportions of juveniles and adults (denoting population age structure) are shown in rounded percent for two censuses (Day 10 and Day 30). The term 'Pop. size' refers to population size (mean total numbers in the beaker per treatment), as a proxy for reproductive success, is presented to provide a relatively comparative understanding of the proportions. The table also displays the change (increase or decrease) in the proportions on Day 30 compared to Day 10. Ctrl (the control, photoperiod of 24h light:0h darkness, salinity of 0.33 gL$^{-1}$, and pH = ~7), *Salinity1* = 1.33 gL$^{-1}$, *Salinity2* = 3.33 gL$^{-1}$, and *Salinity3* = 6.33 gL$^{-1}$; *Acidity1* (pH = 6), *Acidity2* (pH = 5.5), and *Acidity3* (pH = 5). Sal = Salinity, Acid = Acidity, Sal & Acid = Salinity & Acidity, Photo.p. = Photoperiod, photoperiod in constant light (24h day:0h night), Pop. = Population. *Daphnia* age structure refers to the proportions of juveniles relative to the total beaker population per treatment combination.

| | | Day 10 | | | Day 30 | | | Increase/decrease on Day 30 compared to Day 10 |
|---|---|---|---|---|---|---|---|---|
| **Photoperiod** | **Treatment** | **Pop. size** | **% Juveniles** | **% Adults** | **Pop. size** | **% Juveniles** | **% Adults** | **Adults** |
| 24D:0N | Ctrl | 16 | 65 | 35 | 328 | 98 | 2 | 33 % less |
| 24D:0N | *Sal1* | 42 | 88 | 12 | 102 | 84 | 16 | 4 % **more** |
| 24D:0N | *Sal2* | 10 | 54 | 46 | 110 | 80 | 20 | 26 % less |
| 24D:0N | *Sal3* | 27 | 74 | 26 | 271 | 91 | 9 | 17 % less |
| 24D:0N | *Acid1* | 19 | 56 | 44 | 52 | 84 | 16 | 28 % less |
| 24D:0N | *Acid2* | 7 | 37 | 63 | 131 | 97 | 3 | 60 % less |
| 24D:0N | *Acid3* | 5 | 33 | 67 | 57 | 100 | 0 | 67 % less |
| 24D:0N | *Sal1 & Acid1* | 32 | 79 | 21 | 97 | 64 | 36 | 15 % **more** |
| 24D:0N | *Sal2 & Acid2* | 12 | 54 | 46 | 292 | 84 | 16 | 30 % less |
| 24D:0N | *Sal3 & Acid3* | 16 | 66 | 34 | 67 | 90 | 10 | 24 % less |

*Note1: Daphnia population size and age structure differences in salinity, acidity, and their combination under constant darkness (24h)*

We conducted a follow-up to the primary experiment described in the main text, where photoperiod was lacking in terms of constant absence of light (24h dark). We repeated the 9 stress combinations (*Salinity1* = 1.33 gL$^{-1}$, *Salinity2* = 3.33 gL$^{-1}$, and *Salinity3* = 6.33 gL$^{-1}$; *Acidity1* [pH = 6], *Acidity2* [pH = 5.5], and *Acidity3* [pH = 5]. Sal = Salinity, Acid = Acidity, Sal & Acid = Salinity & Acidity) plus a control (the standard ADaM, as specified in the main-text Methods, with optimal conditions [photoperiod of 16h light:8h darkness, salinity of 0.33 gL$^{-1}$, and pH = ~7]), with 7 early instars in the beaker, and 3 replicates per treatment combination. This resulted in 196 daphnids compartmentalised into the combinations under the 24h darkness conditions. Population size (total number of daphnids in the beaker at each census, as a proxy of reproductive success) and the proportion of juveniles relative to the total population in the beaker (age structure, hereafter) were examined after 30 days of the *Daphnia* introduction. To ensure complete darkness, each beaker was completely double-wrapped with foil, with an obscured sealable opening for feeding with pipette every other day. The media were not changed during the course of this part of the work to ensure the extremity of the limiting or adverse environmental conditions; data collection took place only at Day 30 because we could not count the *Daphnia* without exposing them to light and it was essential to eliminate all exposure to light and maximise the adversity of environmental challenge during this part of the work. We addressed the following question: What are the changes in *Daphnia* traits (population size and age structure) in response to the effects of salinity and/or acidity under prolonged continuous darkness?

Exposure to the physico-chemical stress levels in the dark, caused all but very few individuals to survive these extreme conditions to the 30-Day census. As such, we provide descriptive statistics only. We made three major observations: First, the only treatment in the 24-hour dark phase where individuals in all three replicates survived was Salinity1; the average population increased by only 43% at Day 30 in comparison with Day 1, where juveniles made up the majority (84%) of the population, Supplementary Information (Table S9). Second, the population of one replicate only was retrievable under *Acidity1* but the population was extremely small and contained no adult individuals marking a decrease of 95% from the initial count status on Day 1. And third, the population under combined stress *Salinity1 & Acidity1* was 48% larger on Day 30 and had similar proportions of adults and juveniles, Supplementary Information (Table S9). These observations suggest that the lack of photoperiod had a significant negative impact on the ability to tolerate and survive chemical stressors in *D. magna* and that the survival in the darkness was associated with a specific single level of salinity, acidity, and their combination, hinting at certain thresholds beyond which *D. magna* seemed to fail to survive in the dark. Notably, the ADaM medium in this experiment did not show any signs of foulness or turbidity in any of the beakers. This suggests that there were no major negative effects of decaying organic material, due to the salinity of the medium, behind the observed high mortality after the 30 day-spell of constant darkness. However, the *intended* lack of water exchange, mimicking entrapment in water in cavities or underground with limited exchange, accumulation of catabolic substances, and gradual shortage of nutrients per capita, might have led to poorer water quality that could explain the extremely low survival in the dark. Nevertheless, our results imply that specific levels of chemical stress might have contributed to the ability of small parts of the population to survive as all daphnids failed to survive constant darkness when it was the only stressor applied. Our findings receive support from Connelly et al. (2016) asserting that poor or abnormal light conditions can compromise *Daphnia* survival[1]. Furthermore, amid the variety of adverse effects of freshwater acidification[2,3], disruption of olfactory and chemosensory perception, which are crucial especially under low light conditions, has been shown to result in maladaptive response to conspecific and heterospecific cues by aquatic organisms[4,5]. Lastly, a deductive reasoning of the general outcomes of chemical stress under constant light and constant darkness clearly indicates a far more negative impact of the constant absence of light on *D. magna* populations with differential effects on age structure. See also Supplementary Information *Note 2*.

**Table S9. *Daphnia population* size and age structure differences in salinity, acidity, and their combination under constant darkness (24h).** Mean population sizes and rounded proportions of juveniles and adults are shown for Day 30, corresponding to the treatments *Salinity1*, *Acidity1*, and their combination *Salinty1 & Acidity1*. Their treatment counterparts, under constant light, plus a control are displayed for a general comparative purpose. Ctrl (the control, photoperiod of 16h light:8h darkness, salinity of 0.33 $gL^{-1}$, and pH = ~7), *Salinity1* = 1.33 $gL^{-1}$, *Salinity2* = 3.33 $gL^{-1}$, and *Salinity3* = 6.33 $gL^{-1}$; *Acidity1* (pH = 6), *Acidity2* (pH = 5.5), and *Acidity3* (pH = 5). Sal = Salinity, Acid = Acidity, Sal & Acid = Salinity & Acidity, photoperiod in constant darkness (0h light:24h night), photoperiod in constant light (24h day:0h night). *Daphnia* population size refers to the total numbers in the beaker per treatment combination, *Daphnia* age structure refers to the proportions of juveniles relative to the total beaker population per treatment combination.

| Photoperiod | Treatment | Population size on day 30 | % Juveniles | % Adults |
|---|---|---|---|---|
| 16D:8N | Ctrl | 119 | 92 | 8 |
| **0D:24N** | *Sal1* | 10 ±6.94 SEM | 19 | 56 |
| **0D:24N** | *Acid1* | 0.33 ±0.27 SEM | 100 | 0 |
| **0D:24N** | *Sal1 & Acid1* | 10.33 ±8.44 SEM | 52 | 48 |

*Note 2: Remarks on the relevance of the study's design and findings with regard to the multifaceted environmental challenge of today's and tomorrow's worlds*

The primary focus of the current study is to trigger and examine phenotypic plasticity responses (reproductive and developmental) in *Daphnia magna* as an important biological indicator, model organism, and fish feed under severe complex conditions. Given the dearth of study on the effects of composite chemical stress under the lack of photoperiod, our work seeks to fill some gaps against this background and suggests future directions for an investigation of differential phenotypic expressions in response to challenging environmental contexts. Our findings highlight *Daphnia,* a keystone species, as a promising model system for evolutionary studies on phenotypic plasticity and epigenetics[6,7], especially in the sphere of examining combined effects of multiple environmental stressors[8] in novel or atypical environments.

Anthropogenic illumination of more areas at night may bring civil and economical benefits[9]. However, it also comes with a hefty price in terms of altering the biological rhythm of water life plus to greenhouse gas emission that eventually contributes to the aggravation of climate change and precipitation, all together resulting in more acidification and salinisation of freshwater habitats[10,11]. Literature review on artificial light at night suggests that ecological light pollution accompanied by changes in water quality and conditions in lentic and lotic systems[11-13] may lead to significant consequent changes in species physiology[14]; population sizes, composition, richness, ranges and distribution of species[11-14]; heightened prey capture due to enhanced predator vision[13] and increased foraging[15] with species-dependent positive or negative phototactic responses[15]; modification of food-webs due to behavioural changes in zooplankton feeding on algae[11,16] and thus leading to considerable ecological and micro-evolutionary changes[15,17]. Concomitant with alteration of ecoevolutionary dynamics, feedback and local adaptations of species subject to complex selective pressures in the vulnerable systems[18,19], the collection of these effects and changes is destined to lead to alteration of ecosystem functioning[11,13]. The lingering question, however, is about how the lack of photoperiod, *e.g.*, constant light, may underlie plastic daphnid responses and by which mechanisms; a matter that lays the basis for much needed further molecular investigation.

The disturbance of nighttime underwater illumination can interfere with activities associated with lunar light, and may lead to upsetting of diel vertical behaviour[16], social communication and synchronicity[20,21], reproductive timing and sync[9,20,21], and predator-prey interactions for aquatic life forms including zooplankton[9]. For instance, Kaniewska et al. (2015) demonstrated that light pollution related to urbanisation created a state of discrepancy in cellular signalling that led to precluding spawning in corals due to out-of-sync biological time[20]. We may, therefore, speculate that the extended light exposure would lead to alteration of organism response to other chemical stressors that can boost or lessen their chances to survive adversity. Changing nighttime pattern in *Daphnia* because of artificial lighting has been documented to disrupt their floating to capture algal food as a result of altering their diel vertical migration[16], which in turn can result in algal blooms that challenge other life forms through changing water quality and obstructing sunlight in ponds and lakes[16,17,22]. This can get aggravated, as well, by intrusive anthropogenic alteration of the ecosystem[12,23]. The interaction between chemical stressors and the disruption or masking of circadian rhythmicity[9,21], due to extended light or darkness exposure, is indeed an important aspect of the multi-faced environmental challenge of the world of today that has not been adequately investigated and delineated well thus far[11,13]. The species or genotypes within species that can locally acclimate and subsequently adapt to those complex freshwater selective pressures are expected to have an evolutionary advantage and perhaps a degree of invasiveness of the novel habitats[19,24], as other taxa that fail to adapt may lose to invasive and exotic species that can tolerate the combined stress fallout[12,24,25].

The effects of altered photoperiod on zooplankton survival in acidified water are of paramount importance as under decreasing pH, *Daphnia* would become displaced by other arthropods[24,25]. A similar outcome can be pictured when abnormal light exposure is accompanied by acute salinisation beyond *Daphnia* halotolerance capacity to the benefit of halophiles. The effects and impacts of physical and chemical stress discussed above are predicted to intensify/ramify when accompanied with combined chemical stress for example increased mortality of when spending more time in light and chemically polluted areas[13,26-29]. It should be noted that the ability of *D. magna* to regulate osmolarity in the face of external ionic challenge is a result of the remarkable capacity of this organism to counterbalance the effects of salinity beyond 5 $gL^{-1}$ by switching from osmoregulation to osmoconformance which enables the daphnids to cope with increased salinisation[30-31]. For example, the findings reported in the main text of our present study clearly demonstrate that the largest population sizes were generally under osmoregulation in the short run (Day 10); that was the case, as well, in the long run for combined stress *Salinity2 & Acidity2* (Salinity2 = 3.33 $gL^{-1}$ and Acidity2 [pH = 5.5]); but then there was a shift to osmoconformance in *Salinity3* (6.33 $gL^{-1}$), which increased survival and reproduction in the longer term (Day 30). Furthermore, over time *D. magna* may acclimate to the upper boundary of salinity (halotolerance), just below lethal levels contingent on the acidity of the system. Such effects might have occured within the 30-day period of our study.

All individuals used in our experiments were genetically identical as they came from a single mother and comprised the stock culture of the parthenogenetic clone. It is plausible, therefore, that the variable population dynamics and age structure across treatments can be characterised by within-clone ecological differentiation[32-34] arising from stress-induced phenotypic plasticity of the particular *D. magna* genotype used here[33], but see evidence provided by Jose et al. (2009) who found only small differences in the metabolic capacity across different clones of *Daphnia pulex* in response to pH and thermal treatments[34]. We, nevertheless, speculate that different clones with different life histories or communities of mixed clones, or colonies with sexual morphs, would show differential responses as governed by the contexts they are embedded in, their tolerance, and their life histories[34-36]. Our study and findings also provide an example of artificial selection brought about by man that can maximise pressure on the organism to acclimate and adapt or dwindle away and perish, especially in our continuously urbanising world[37]. Elevated artificial light exposure can modify and interfere with natural light-mediated processes across all levels from individual to population to community to ecosystem[9,11,21] and with the expansion of human population the impacts of artificial light at night on trophic interactions and ecosystems are expected to be more intensive and extensive in the future[11].

The model ecosystem of our study finds relevance and provide implications for future *in-situ* studies under scenarios where light and water physico-chemical conditions are subject to numerous primary and secondary alteration or disruption: Volcanic aftermath through the complex tsunami-triggering as well as physico-chemical environment-altering effects including reflecting, scattering and obscuring of sunlight (*e.g.,* floating pumice on lake surfaces) caused by volcanic tephra emissions and debris spewed into the air, soil, and waters[38-41]; vog (volcanic smog)[42,43]; clouds, haze, fog, and aerosol particles[44-46]; air-borne pollution and smog[47,48]; water turbidity[49]; deforestation as well as alteration of natural and forest canopy and riparian vegetation[49-53], forest fire smoke[54-56]; ground water exposure and pond water-tunnelling underground through natural and human-induced sinkholes and cenotes resulting in acidified water from rock dissolution[57-59]; the dark of bromeliad water-retaining phytotelmata[60,61]; urbanisation[9,37]; aquaculture farming and fishing under artificial light conditions[10,17,62]; algal blooms/scum/mats[12,22,63,64] that are elevated by agriculture[23]; and periphyton cover[50,51] as increasing production and hence artificial shade of periphyton is associated with decline in total invertebrate density and water quality[64] in freshwater systems[50-52].

Moreover, there is an important economic dimension of our study because our results suggest that the absence of photoperiod (in terms of constant light) enhances and improves the tolerance of *Daphnia* (zooplanktonic food for fish), as well the daphnid fecundity and plasticity against chemical stress. It is a case where a stressor is jolting or boosting the system of the organism to perform better and hyper-reproduce under escalated chemical stress. As such, our study findings provide promising implications for fish farming through boosting the osmoregulatory capacity, tolerance, and reproduction of zooplankton under constant light for specific periods prior to releasing into aquaculture farms. It is worth mentioning that fishing with light is increasingly popular for commercial and industrial reasons, albeit being widely favourable and profitable this method has considerable hazards for the sustainability and well-being of aquatic life and fisheries [5 >>>10]. It yet remains to be studied, however, whether the effect of increased light exposure at night on enhancing the reproductive plasticity may be universal for other freshwater taxa.

Last but not least, our findings highlight the utility of *D. magna* as an extremely plastic model organism in next-generation biotechnology studies under severe conditions[65]. This can also be utilisable in assays where constant darkness, or especially constant light, triggering extreme osmoregulation and alteration of body clock, may prevail for considerable spells of time as in orbital stations[66,67]. Investigation of extremophiles, *i.e.,* organisms that can survive extreme environmental conditions, and testing extremely phenotypically plastic organisms under severe conditions are gathering momentum because of their myriad eco-evo implications and applications. This includes, by no means as an exhaustive list but as good indication of this futuristic much needed stream in science, testing the survival of microorganisms in extreme environments[68], Tardigrades (water bears) versus radiation[66], and resurrection ecology[69]along with revitalisation from the verge of death [32] in *Daphnia*. The 'extremophilia' as a process (or phenomenon) has been receiving increasing attention mainly in terms of thermal and geochemical stress conditions, but the investigation of extremely plastic organisms and extremophiles under the extreme absence of circadian rhythm/photoperiod accompanied by combined chemical stress (salinisation and acidification) is surprisingly understudied. We wish to stress out that "…extreme environments are usually an anthropogenic construct based on extremes of the physical environment around a baseline…" *sensu* Bell (2012)[70].
